# Superloser: a plasmid shuffling vector for *Saccharomyces cerevisiae* with exceedingly low background

**DOI:** 10.1101/630863

**Authors:** Max A. B. Haase, David M. Truong, Jef D. Boeke

## Abstract

Here we report a new plasmid shuffle vector for forcing budding yeast (*Saccharomyces cerevisiae*) to incorporate a new genetic pathway in place of a native pathway – even essential ones – while maintaining low false positive rates (less than 1 in 10^8^ per cell). This plasmid, dubbed “Superloser”, was designed with reduced sequence similarity to commonly used yeast plasmids (i.e. pRS400 series) to limit recombination, a process that in our experience leads to retention of the yeast gene(s) instead of the desired gene(s). In addition, Superloser utilizes two orthogonal copies of the counter-selectable marker *URA3* to reduce spontaneous 5-fluoroorotic acid resistance. Finally, the CEN/ARS sequence is fused to the *GAL1-10* promoter, which disrupts plasmid segregation in the presence of the sugar galactose, causing Superloser to rapidly be removed from a population of cells. We show one proof of concept shuffling experiment: swapping yeast’s core histones out for their human counterparts. Superloser is especially useful for forcing yeast to use highly unfavorable genes, such as human histones, as it enables plating a large number of cells (1.4×10^9^) on a single 10 cm petri dish while maintaining a very low background. Therefore, Superloser is a useful tool for yeast geneticists to effectively shuffle low viability genes and/or pathways in yeast that may arise in as low as 1 in 10^8^ cells.

---

The plasmid shuffling method in *Saccharomyces cerevisiae* facilitated the straightforward genetic study of essential genes and pathways by enabling the replacement of wildtype alleles for either temperature sensitive mutants or distant homologs (Boeke *et al.* 1987; Budd and Campbell 1987; Mann *et al.* 1987; Sikorski and Boeke 1991; Forsburg 2001). This method works by having a null allele of a gene on the chromosome, kept alive by a wildtype copy on a counter-selectable plasmid - typically employing the *URA3* marker. Next, a second plasmid with a mutant allele and a different auxotrophic marker is added and the two plasmids are exchanged (“shuffled”). Counter-selection of the wildtype allele on the *URA3* plasmid using the compound 5-Fluoroorotic acid (5-FOA), which is toxic to *URA3*^+^ cells but not to *ura3* mutants, selects for loss of the wildtype gene in favor of the mutant allele (Boeke *et al.* 1987). This classic yeast genetics tool is routinely used, but as more adventurous shuffle experiments “on the edge” of selective power are attempted a new tool is needed to minimize the impact of false positives associated with plasmid shuffle events.

We recently used plasmid shuffling to show that budding yeast can survive using human core histones in place of their own (Truong and Boeke 2017). However, we found this to be an extremely rare event taking three weeks and occurring in substantially less than 1 in 10^7^ cells. As a result, counterselection during plasmid shuffling favors the formation of background colonies, making it difficult to identify true shuffle events. These background colonies consist of two major types: 1) spontaneous inactivation of the *URA3* marker 2) or formation of plasmid recombinants in which the incoming plasmid which lacks *URA3* acquires the essential gene (e.g. yeast histones), or the resident plasmid acquires the incoming selectable marker. One factor promoting formation of the recombinants is the regions of sequence identity shared between the commonly used yeast shuffle vectors of the pRS400 series (Sikorski and Hieter 1989). Because of this, the ratio of true positives to false positives is considerably lower therefore decreasing the likelihood of isolating correct colonies. As yeast has become a favored organism for industrial biotechnology (Buijs *et al.* 2013; Paddon *et al.* 2013; Mattanovich *et al.* 2014; Galanie *et al.* 2015; Awan *et al.* 2017; Luo *et al.* 2019) new opportunities for complex cellular engineering mandate more robust tools for gene/pathway swapping.

Here, we developed a new plasmid shuffling vector dubbed the “Superloser plasmid” - alluding to its inability to survive 5-FOA counterselection, resulting in exceedingly low background when used in plasmid shuffling experiments. The Superloser plasmid has very short (and strategically inverted) regions of sequence identity to commonly used yeast vectors, and its loss from the population can be promoted by galactose induction, which drives transcriptional inactivation of the centromere. The combined effects of these features is a system with a much lower background of 5-FOA^R^ variants in shuffling experiments. We show that Superloser lowers the background of the yeast histone humanization plasmid shuffle by >99% compared to use of the commonly used *URA3* plasmid pRS416, and increased the frequency of observing true positives by 13-fold. These results demonstrate that Superloser is a useful tool for yeast geneticists and demonstrates that it is ideal in situations where the genes to be swapped cause a severe decrease in cellular fitness, but can also be readily used for manipulation of any gene or pathway.

## Materials and Methods

### Strains and media

All yeast strains used in this study are described in Table 1. 5-Fluoroorotic Acid Monohydrate (FOA, 5-FOA) was purchased from US Biological (Cat. F5050) and used at a concentration of 1 mG mL^−1^. Yeast strains were cultured in yeast extract peptone dextrose (YPD) or synthetic complete media (SC) with appropriate amino acids dropped out. All yeast transformations were carried out using standard lithium acetate procedures.

**Table 1.**
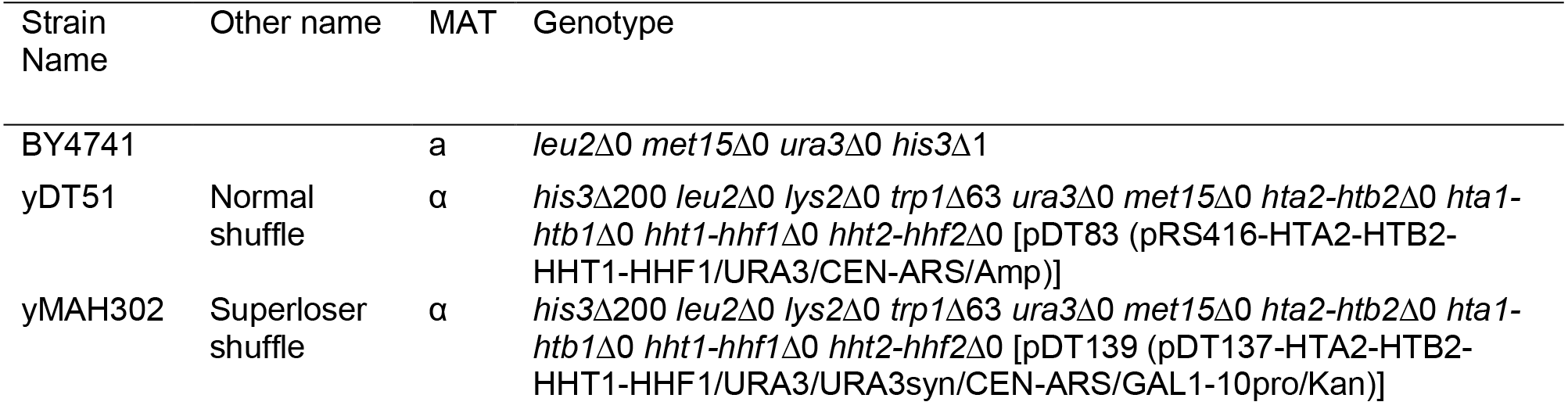
Strains used in this study.

Cloning was performed in Top10 *Escherichia coli* grown in either Luria Broth (LB) or Super Optimal broth with Catabolite repression (SOC) media. To select transformants with drug resistant genes, either carbenicillin (75 μg mL^−1^) or kanamycin (50 μg mL^−1^) was used where appropriate.

### Plasmids and oligos

All plasmids used are listed in Table 2, oligos used are listed in Table 3. We derived the backbone of Superloser v1 (pDT137) from pTwist30 (Twist Biosciences). Relevant genetic parts were amplified by PCR and assembled using Gibson assembly (Gibson *et al.* 2009). Additionally, the *GAL1-10* promoter was placed adjacent to the CEN/ARS sequence - with the *GAL1* promoter oriented towards CEN ensuring transcription at the *GAL1* promoter will destabilize the centromere. To construct Superloser v2 (pMAX175), LacZα was PCR amplified from pUC19 and Gibson assembled with PCR amplified fragments of *URA3*, *URA3*syn, and the Superloser v1 backbone. Both Superloser v1 and v2 have been deposited with Addgene and are available upon request.

**Table 2.** Plasmids used in this study.

| Plasmid name | Other name | Plasmid markers | Description |
| --- | --- | --- | --- |
| pDT83 | yHistones | *amp/URA3* | pRS416-HTA2-HTB2-HHT1-HHF1. Shuffle plasmid, parental strain |
| pDT109 | hHistones3.1 | *amp/TRP1* | pRS414-hH3.1-hH4-hH2A-hH2B (HHT2F2HTA1B1 PROs/TERs). |
| pDT137 | Superloser-v1 | *kan/URA3* | pTwist30-GAL1-pro/CEN-ARS/URA3/URA3syn |
| pDT139 | Superloser yHistones | *kan/URA3* | pDT137-HTA2-HTB2-HHT1-HHF1/URA3/URA3syn/CEN-ARS/GAL1-10pro/Kan) |
| pMAX116 | pRS416-LEU2 | *amp/URA3/LEU2* | pRS416-LEU2 |
| pMAX121 | Superloser-LEU2 | *kan/URA3/LEU2* | pDT137-LEU2 |
| pMAX175 | Superloser-v2 | *kan/URA3* | pTwist30-GAL1-pro/CEN-ARS/URA3/LacZα/URA3syn |

**Table 3.**
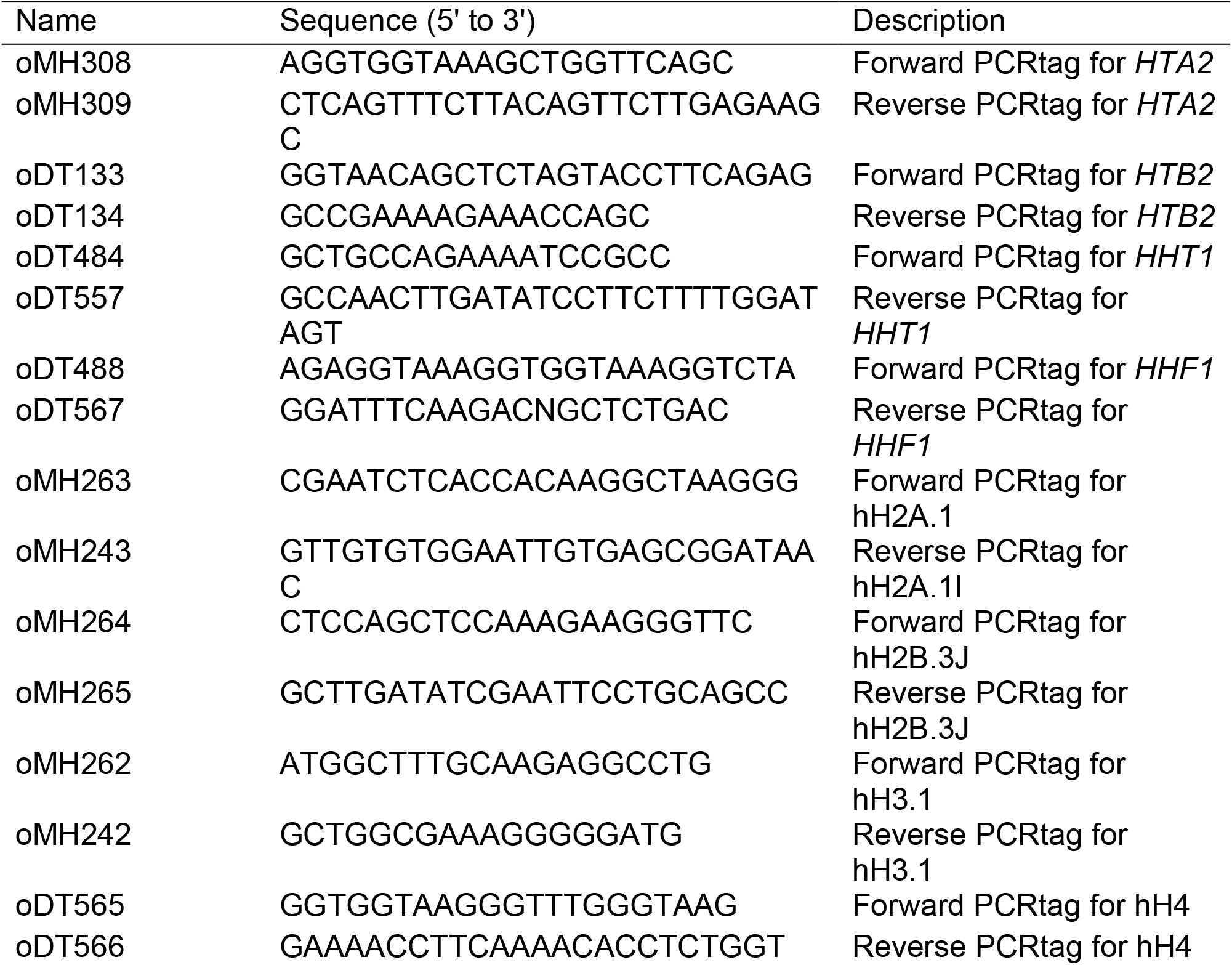
Oligos used in this study.

### Spontaneous 5-FOA resistant mutant assay

BY4741 was transformed with either pDT137-*LEU2* or pRS416-*LEU2*. Forty-six transformants from each were grown overnight in 1 mL of SC-Leu-Ura in a 96-deep well plate. The following day, the cultures were centrifuged at 3000 rpm for 3 min to remove the medium, washed with sterile water, and resuspended in 1 mL of SC-Leu medium. These cultures were then passaged daily for either one or five days in SC-Leu. On the final day, the cultures were centrifuged as before and resuspended in 100 μL of sterile water. From the resuspension, 10 µL was removed and micropipetted onto SC-Leu+5-FOA agar plates (in four replicates). After one week, the number of spontaneous 5-FOA^R^ mutants was determined by colony counting.

To calculate the mutation rates, we utilized the *P*_0_ method (Lea and Coulson 1949). Briefly, the mean number of mutations is calculated by taking the zeroth term of the Poisson distribution (the number of cultures with no mutants). Next, the mean number of mutations is calculated as the −ln(*P*_0_) and this value is divided by the number of cells dotted out to give the mutation rate.

### Galactose induced plasmid destabilization

An additional means of reducing background is to directly encourage plasmid loss by introducing a *GAL* promoter next to the centromere sequence on the plasmid. The plasmid is then predicted to become much less stable on galactose medium (Murray and Szostak 1983; Panzeri *et al.* 1984; Chlebowicz-Sledziewska and Sledziewski 1985; Hill and Bloom 1987a; Beach *et al.* 2017). To verify that this destabilization was occurring in the Superloser we co-transformed BY4741 with either pRS415/pRS416 or pRS415/pDT137. Three clones were picked and grown overnight in SC-Leu-Ura. The following day, the cultures were washed with sterile water, resuspended in 5 mL of sterile water, and 50 µL was used to inoculate 5 mL of SC+2%dextrose and SC +2%galactose/1%raffinose. Additionally, the washed and resuspended cells were serially diluted and plated on YPD (approximately 250 cells per plate). The cultures were then passaged for three days, and each day ~250 cells were plated to YPD agar. Finally, each YPD plate was replica-plated to both SC-Ura and SC-Leu agar plates and the number of colonies retaining each plasmid was determined. Rate of loss for each plasmid was calculated as follows: First, semi-log nonlinear regressions were performed on the data from each biological replicate of each plasmid (n=3). Second, using the regressions the mean percent loss per cell per generation was calculated as the percent change between timepoints divided by the number of generations per day (assuming 1 generation per ninety minutes; or 16 generations a day (Sherman 1991)). Lastly, the significance in rate difference between galactose and dextrose for each plasmid was determined by one-way ANOVA analysis of the calculated rates.

### Yeast to human histone plasmid shuffle

Histone shuffle strain yDT51 (Truong and Boeke 2017) which contains a set of yeast histones on a standard *URA3 CEN* vector, was transformed with a *TRP1* plasmid containing the human histones. In addition, a second shuffle strain (yMAH302), which contains a set of yeast histone genes on the Superloser plasmid, was transformed with the same *TRP1* plasmid containing the core human histone genes. Colonies were selected at 30°C on SC-Ura-Trp plates. 10 colonies of each transformation were picked and grown overnight in both SC-Trp+2%dex and SC-Trp+2%gal+1%raf. The next day, the OD600 of each culture was taken and the entirety of each culture was plated onto SC-Trp+5-FOA plates. Plates were then incubated at 30°C in a sealed plastic tupperware container for up to four weeks with a napkin wetted with sterile water for humidification. Colonies were counted weekly and any colonies appearing after 3 weeks were analyzed by PCRtag analysis (Mitchell *et al.* 2015; Truong and Boeke 2017) to confirm humanization. Primers used for PCRtag analysis are listed in Table 3. GoTaq Green Master Mix (Promega, WI) was used for amplification with the following PCR conditions were used: 35 cycles of 98° for 30sec, 55° for 30sec, and 72° for 45sec.

## Results

### Design of the Superloser Plasmid

We designed Superloser to be maximally effective for “aggressive” plasmid shuffle experiments of essential genes, such as simultaneous interspecies swapping of all four of the core histones (Truong and Boeke 2017). To accomplish this, we designed in three features we reasoned would reduce the background in plasmid shuffling experiments: (1) markedly reduced sequence identity to commonly used yeast plasmids, (2) an inducible mechanism to promote rapid loss of the plasmid (3) and redundant, but orthogonal, *URA3* markers for 5-FOA counterselection. In order to reduce sequence identity we derived the backbone of Superloser from pTwist30, which has only a minimal pUC origin and a KanR marker with little intervening sequence. To promote plasmid loss we fused the *CEN/ARS* sequence from pRS400 to the *GAL1-10* promoter (*GAL1p*-CEN), with the *GAL1* promoter orientated to transcribe though the CEN region; a strategy successfully used for inducing chromosomal aneuploidies (Hill and Bloom 1987b; Beach *et al.* 2017). We then designed in a second orthogonal *URA3* marker (*URA3*syn), which was recoded for maximally different codon usage and is controlled by an orthogonal promoter from *MET17* and terminator from *HIS3*.

In addition, we constructed a second version of Superloser that contains the common *LacZ*α cloning site (Figure 1B 1C). Importantly, we ensured the native *URA3* gene and *LacZ*α gene and multiple cloning site were inversely oriented to the *CEN/ARS* sequence in comparison to the pRS400 series of vectors. In order to test the utility of Superloser we performed a series experiments to see if each designed feature worked as intended and if each feature contributed to lower background of plasmid shuffling.

**Figure 1.**
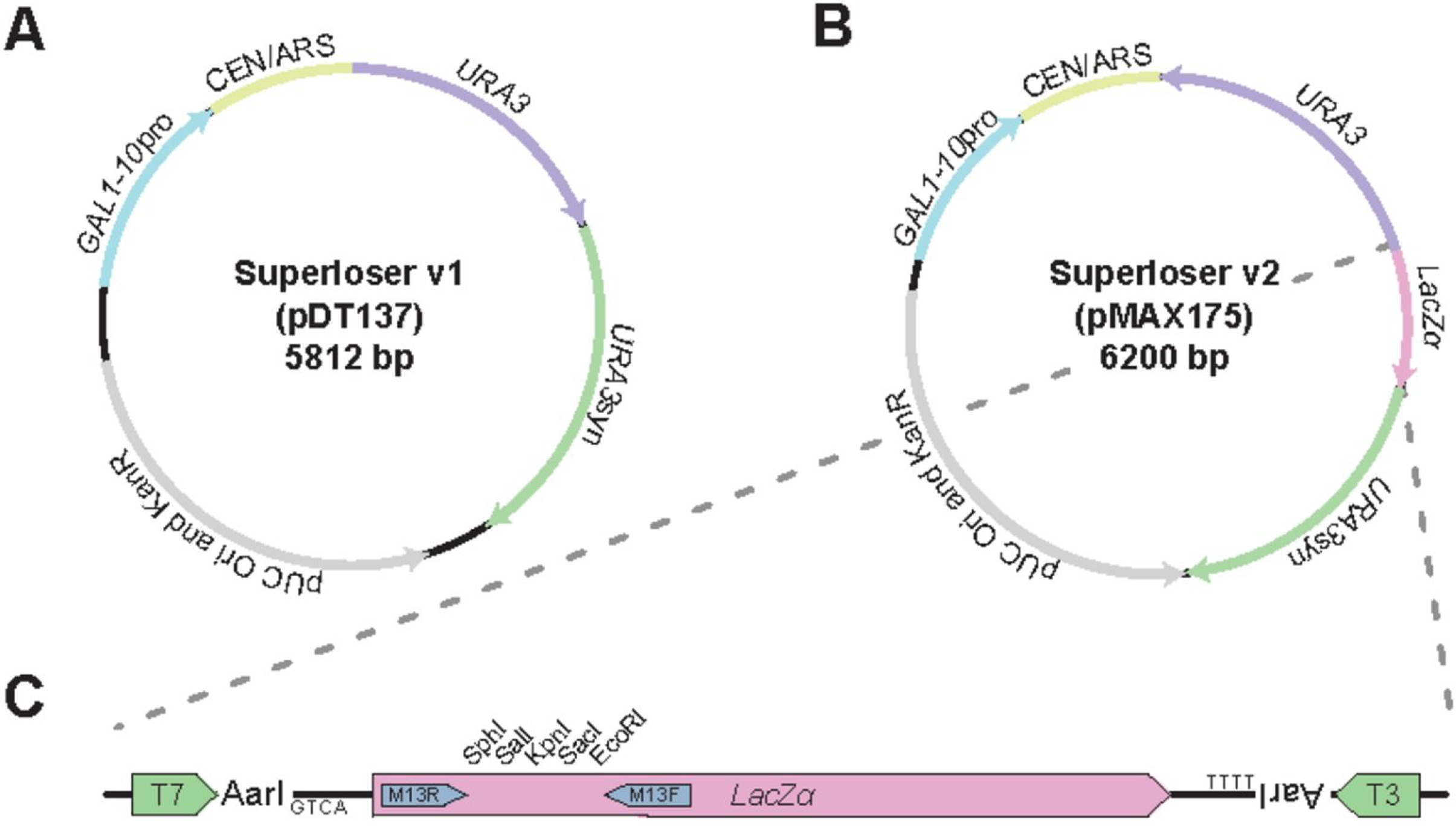
Design of the Superloser plasmid. (**a**) plasmid map of the Superloser version 1, which was used in this study. (**b**) Superloser version 2 contains a *lacZα* cloning site enabling blue-white colony screening. (**c**) Zoomed view of the unique restriction sites of Superloser v2.

### Dual *URA3* markers reduce background 5-FOA resistance

We first asked if the addition of the *URA3*syn marker decreased the frequency of spontaneous 5-FOA resistance (5-FOA^R^) in Superloser compared to pRS416. To ensure plasmid retention, we cloned in a *LEU2* marker into both pRS416 and Superloser (pDT137). Briefly, 46 clones of BY4741 transformed with either pRS416-*LEU2* or Superloser-*LEU2* were grown overnight in SC-Leu-Ura and the following day overnight cultures were diluted 1:10 and 50 uL was used to inoculate 1 mL of SC-Leu. Cultures were then passaged for 5 days, each day 50 uL of the previous overnight culture was used to inoculate fresh SC-Leu. After the fifth day, a portion of each culture was micropipetted to SC-Leu+5-FOA agar plates to identify 5-FOA^R^ events (Figure 2A, B). In order to estimate the number of cells in each spot, a portion of each culture was pooled, serial diluted, and then plated to YPD agar plates and colonies were counted (average of 8e6 cells per spot). Growth on 5-FOA was monitored for one week and the number of colonies arising in each spot was counted (Figure 3B).

**Figure 2.**
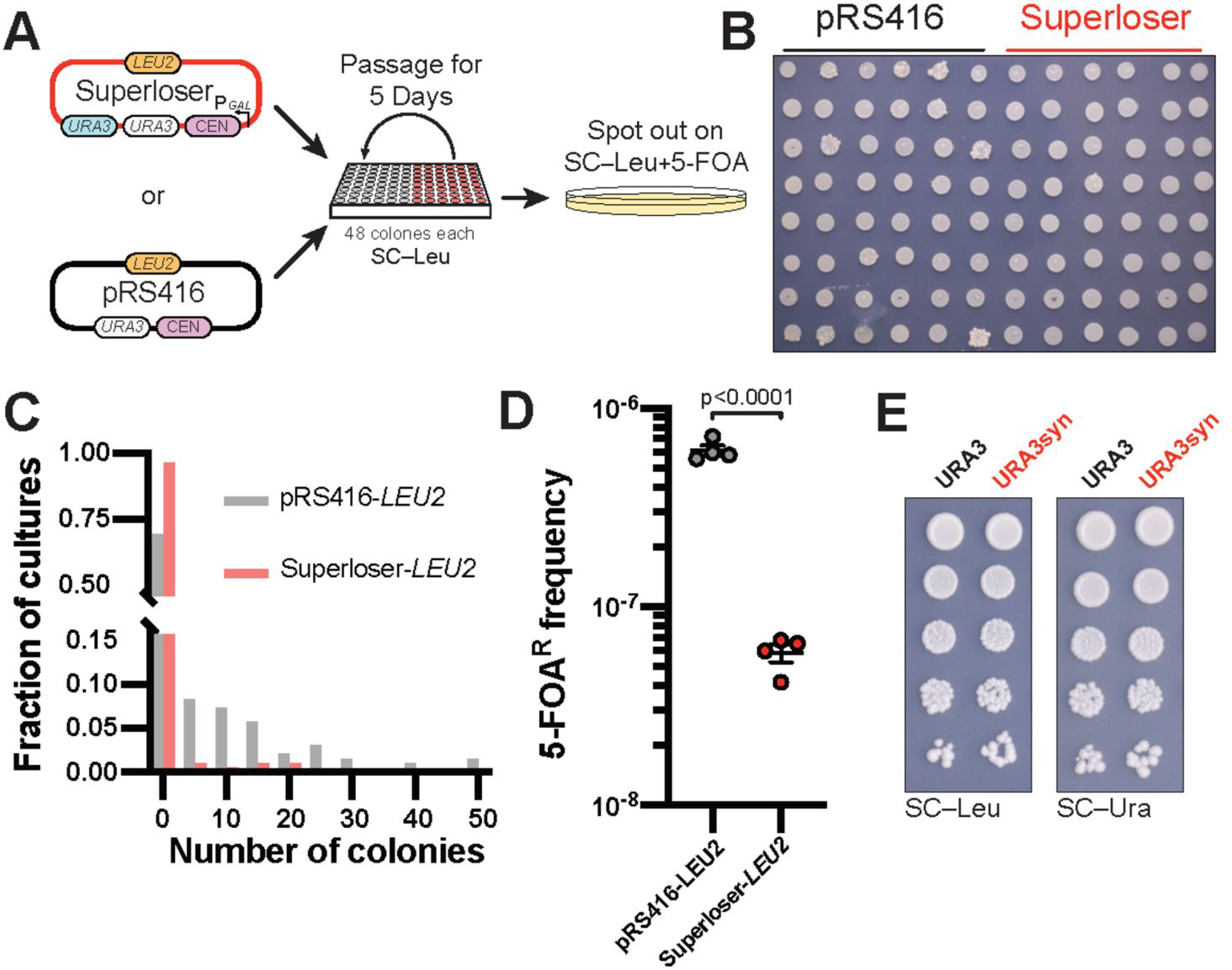
Superloser is robust against spontaneous formation of *ura3* mutants. (**a**) Schematic of experimental setup. Transformants of both Superloser (pDT137-*LEU2*) and pRS416-*LEU2* were passaged for 5 days with no selection of *URA3* and then micropipetted onto SC-LEU+5-FOA trays (see methods). (**b**) Example image of one 5-FOA plate showing 5-FOA^R^ mutants from pRS416-*LEU2* and Superloser (pDT137-*LEU2*). (**c**) Histogram of the number of spontaneous 5-FOA^R^ colonies per culture. (**d**) Frequency of spontaneous 5-FOA^R^ colonies of Superloser (pDT137-*LEU2*) and pRS416-*LEU2*, two-tailed unpaired t-test. (**e**) The recoded *URA3*syn complements Δ*URA3*. Cultures were diluted to OD600 of 0.1, 10-fold serial diluted, and finally micropipetted onto SC-Leu and SC-Ura agar plates.

**Figure 3.**
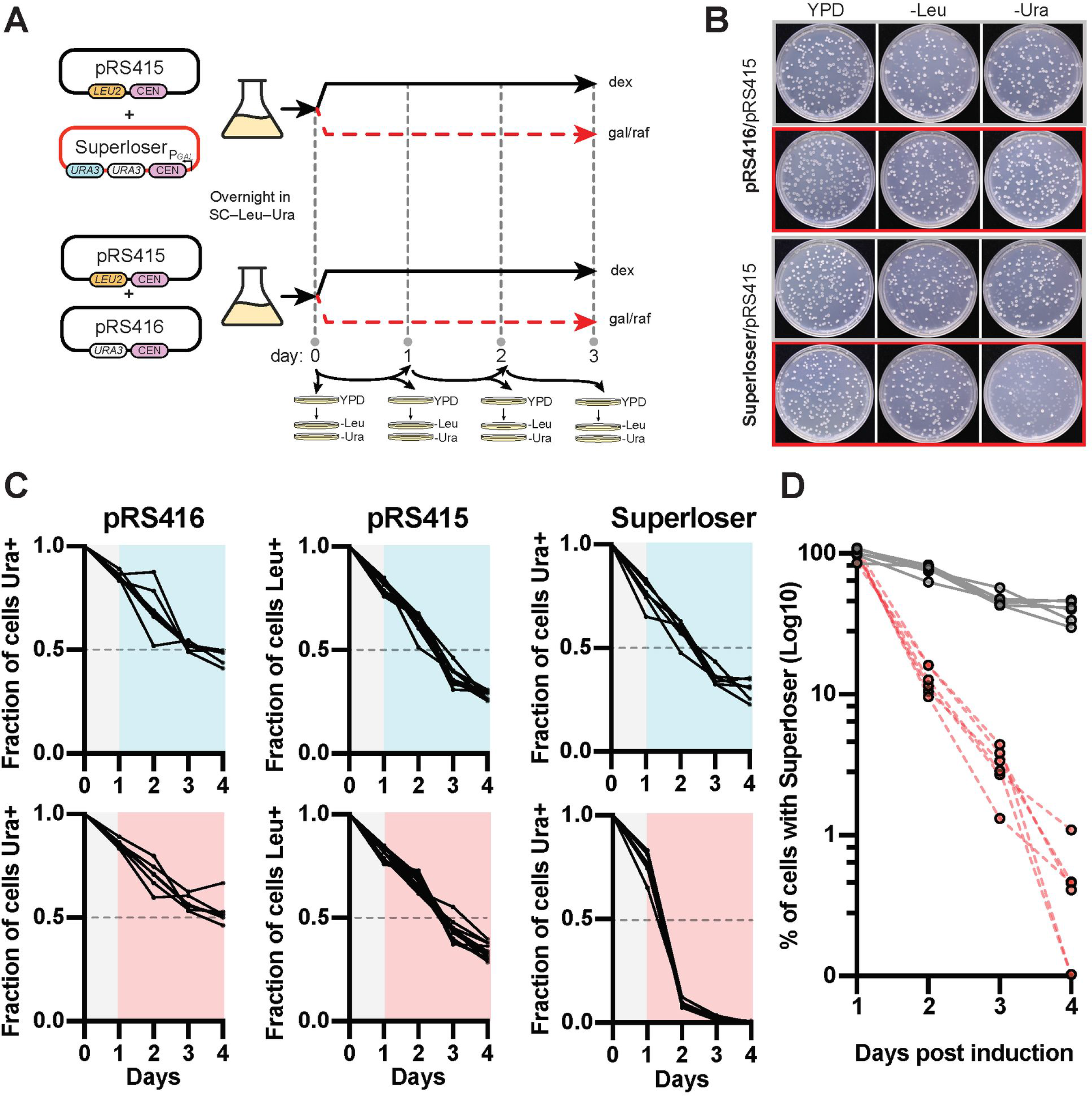
Superloser is rapidly lost in the presence of galactose. (**a**) Schematic of experimental setup. Briefly, three clones of BY4741 co-transformed with the indicated plasmid were cultured overnight in SC-Ura-Leu+2%dex. The following day these cultures were split and sub-cultured into SC complete +dextrose or SC complete +2%galactose/1%raffinose, additionally a portion of each culture were plated to YPD plates in two technical replicates. Each culture was then maintained in the indicated medium with each day a portion of each plated to YPD agar plates in two technical replicates. Finally, each YPD agar plate were replica plated to both SC-Ura and SC-Leu to track plasmid loss. (**b**) Example agar plates after 24 hours growth in SC complete +dextrose or SC complete +2%galactose/1%raffinose. Note the rapid loss of Ura^+^ colonies when in the presence of galactose (red outline, bottom). (**c**) Galactose induced loss of Superloser. The fraction of cells with the indicated plasmid is shown after removal of selection and induction (grey background to colored) under either dextrose (top row, blue) or galactose (bottom row, red). (**d**) Log10 percent of cells with Superloser after induction with galactose or dextrose. Gray dots indicate growth in dextrose, red dots indicate growth in galactose (n=6).

The majority of clones did not give rise to any 5-FOA^R^ colonies for either pRS416 or Superloser, however, Superloser noticeably reduced the number of clones giving rise to more than one 5-FOA^R^ colony (Figure 2C). We next calculated the frequency of 5-FOA^R^ by taking the total number of 5-FOA^R^ colonies observed divided by the number of cells dotted out. Superloser reduced the 5-FOA^R^ frequency by over 10-fold when compared to pRS416 (Figure 2D, student t-test; p<0.0001). Lastly we calculated the mutation rate of both pRS416 and Superloser using the *P*_0_ method, where the mutation rate is estimated from the proportion of cultures with no mutants (*P*_0_) (Lea and Coulson 1949). For pRS416 we calculated a mutation rate of 5.05e^−8^ (8.27e^−9^ - 2.85e^−8^) which is similar to other reported mutation rates of *URA3* (Lang and Murray 2008). Superloser lowered the mutational rate by 19-fold, with a calculated rate of 2.63e^−9^ (1.46e^−8^ – 6.59e^−11^). While the *P*_0_ method is not entirely accurate for *P*_0_ >0.7 (Superloser *P*_0_ = (47/48)), the data presented here shows that the additional *URA3*syn marker in Superloser significantly lowers the spontaneous rate of 5-FOA^R^ mutants.

Lastly, we wanted to confirm that the *URA3*syn marker complemented as well as the native *URA3* marker. To this end, we cloned either the *URA3* or *URA3syn* marker into pRS415 and performed dot assays on SC-Leu and SC-Ura agar plates. We find that the *URA3*syn marker complements Δ*URA3* and is indistinguishable from the *URA3* marker (Figure 2E). In sum, we conclude that the addition of a second *URA3* marker aids in reducing the rate of spontaneous 5-FOA^R^.

### Galactose induction promotes loss of Superloser from the population

We next tested whether the fused *GAL1p-*CEN caused unequal segregation of Superloser plasmid in the presence of galactose. The *GAL1* promoter is transcriptionally inactive in the presence of dextrose, but is rapidly induced upon the switch from dextrose to galactose as a carbon-source (Lohr *et al.* 1995). Transcription through the *GAL1* promoter should disrupt the single point centromeric nucleosome of CEN thereby disrupting kinetochore formation and resulting in the plasmid being retained by the mother cell (Murray and Szostak 1983; Hill and Bloom 1987b; Beach *et al.* 2017). To test whether plasmid missegregation is working as designed we tracked plasmid loss by growth on non-selective media in the presence of either dextrose or galactose (Figure 3A).

As expected, the addition of galactose caused the Superloser plasmid to be rapidly lost from the population (Figure 3B). After 24 hours of induction with galactose only ~12% of cells maintained the Superloser and after 72 hours just ~0.4% kept it (Figure 3B-D). In contrast, in glucose ~77% of cells kept the Superloser after 24 hours and ~39% of cells after 72 hours (Figure 3C, D). Nonlinear regressions were calculated to give the rate of plasmid loss per generation with and without galactose (Table 4). In the presence of dextrose Superloser was naturally lost at a mean rate of 1.6% per cell per generation, similar to the rate of loss of other CEN plasmids (Panzeri *et al.* 1984; Sikorski and Hieter 1989; Baker and Masison 1990). In contrast, in the presence of galactose, Superloser was lost at a mean rate of 5.4% per cell per generation. As controls, we show that loss of either pRS416 or pRS415 was not affected by addition of galactose and each plasmid has similar rates of loss compared to Superloser in dextrose - although we note pRS416 is lost more slowly in comparison to pRS415 or pDT137, perhaps due to its smaller size (Figure 3C and Table 4). Galactose induction therefore effectively and rapidly promotes segregation of the Superloser plasmid from the population - thereby lowering the number of cells retaining both shuffle plasmids upon counterselection and consequently reducing the probability of inter-plasmid recombination.

**Table 4.** Rate of plasmid loss.

| Plasmid | Condition | Mean % loss/cell/<br>generation ±SEM | p value |
| --- | --- | --- | --- |
| pDT137 | Dextrose | 1.6 ± 0.06 | <0.0001 |
| | Galactose | 5.4 ± 0.12 | |
| pRS415 | Dextrose | 1.5 ± 0.06 | 0.7491 |
| | Galactose | 1.4 ± 0.03 | |
| pRS416 | Dextrose | 1.1 ± 0.03 | 0.981 |
| | Galactose | 1 ± 0.05 | |

### Humanization of yeast histones

The benefits of reduced sequence similarity, dual orthogonal *URA3* markers, and inducible loss of Superloser should allow for the shuffling of highly deleterious genes while maintaining low background. As a proof-of-principle for swapping highly unfavored pathways, we revisited the yeast to human histone swap (Truong and Boeke 2017) by comparing the published plasmid system against the Superloser system (Figure 4A). Two isogenic strains harboring the four core yeast histones on either the normal pRS416 (yDT51) or the Superloser (yMAH302) plasmid were transformed with a plasmid containing the four core human histones. Strains were then grown for a day and a half in either SC-Trp+Dex or SC-Trp+Gal+Raf, after which cells were collected and plated onto SC-Trp+5-FOA agar plates. Cells were counted every week for 4 weeks after plating. Any colonies appearing prior to 21 days were considered false positives mostly resulting from interplasmid recombination or inactivation of *URA3* (Truong and Boeke 2017); colonies appearing post 21 days were tested for successful humanization by PCRtag analysis, which distinguishes between highly similar sequences (Mitchell *et al.* 2015; Truong and Boeke 2017).

**Figure 4.**
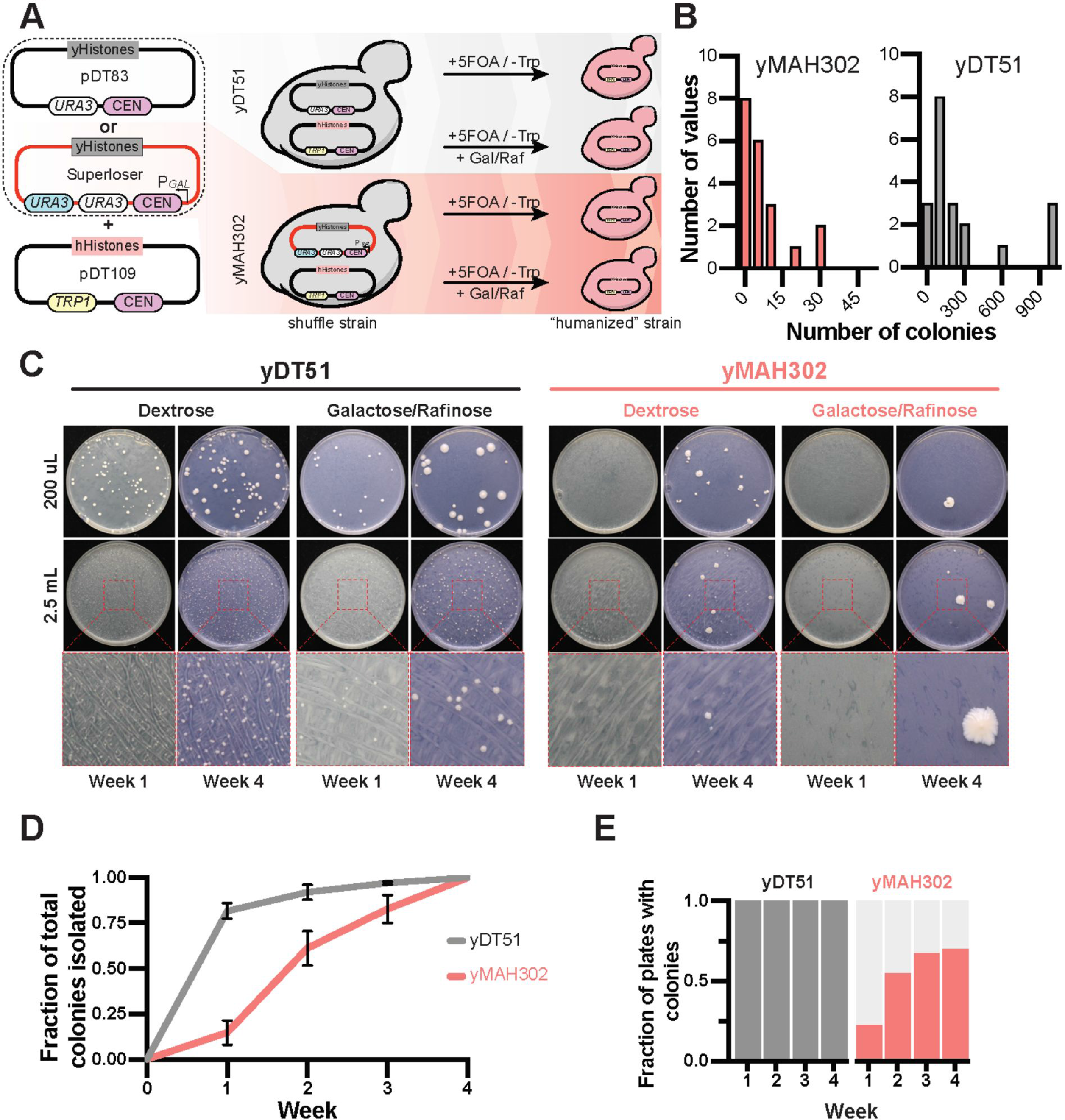
Superloser reduces background associated with humanization of yeast histones. **(a)** Experimental setup: yDT51 with yeast histones on pDT83 (normal) or yMAH302 with yeast histones on pDT139 (Superloser) was transformed with the shuffle plasmid (pDT109) containing the human histones. 20 biological replicates of each strain were grown for 1.5 days in either SC-Trp+2%dex or SC-Trp+2%gal1%raf, at which point cells were collected, washed, and then plated at two concentrations onto SC-Trp+5FOA. Colonies were counted every week for 4 weeks. **(b)** Distribution of the number of 5-FOA^R^ colonies isolated per plate from shuffle experiments with yDT51 and yMAH302. **(c)** Example images of SC-Trp+5-FOA plates from the shuffle experiment. Growth at week 1 is compared to growth at week 4, note the density of colonies at week 1 for yDT51 vs yMAH302. **(d)** Fraction of the total number of 5-FOA^R^ colonies isolated at each week that colonies were observed. Note that most colonies of yDT51 were isolated prior to 3 weeks of growth indicating higher background events. **(e)** The fraction of plates with colonies is shown at each week for yDT51 and yMAH302.

The number of colonies isolated from the normal shuffle strain averaged 277 colonies per plate, whereas from the Superloser shuffle strain averaged 7 colonies per plate - a 38-fold decrease (Figure 4, B and C). This suggested lower rates of recombination between the two shuffle plasmids in the Superloser strain, as expected. Further, we note that the majority of the total colonies isolated from the normal shuffle strain grew prior to three weeks (97%), whereas a smaller portion (82%) grew prior to three weeks from the Superloser shuffle strain (Figure 4D). In addition, all biological replicates from the normal shuffle plasmid gave rise to colonies whereas only ~70% of replicates with Superloser plasmid gave rise to colonies (Figure 4E). Collectively the results strongly indicate decreased background.

After four weeks, the frequency of 5-FOA resistant colonies was calculated as the total number of colonies divided by the total number of cells plated. In dextrose media, the Superloser plasmid resulted in a 26-fold decrease in the frequency of 5-FOA^R^ colonies when compared to the normal plasmid - a 96.1% decrease in the background (Figure 5A; p<0.0001, unpaired t-test). Considering only the Superloser when induced with galactose, we observed a 270-fold decrease in 5-FOA^R^ colonies, or a 99.6% decrease in the background (Figure 5A; p=<0.0001, unpaired t-test). Next, we used PCRtag analysis to test colonies arising post-three weeks for successful humanization (Figure 5B). From the experiments using the Superloser plasmid we isolated 11 successfully humanized clones out of 50 tested. However, we did not isolate any successfully humanized clones when using the normal shuffle plasmid (0/14). In the original study (Truong and Boeke 2017) only 7 such colonies were ever isolated, after very laborious experiments using the conventional shuffle vector. Based on the increased proportion of colonies arising post-three weeks and the dramatic increase in the ease of isolating true positives, we conclude that Superloser effectively shuffles yeast and human histone plasmids while reducing the background tremendously.

**Figure 5.**
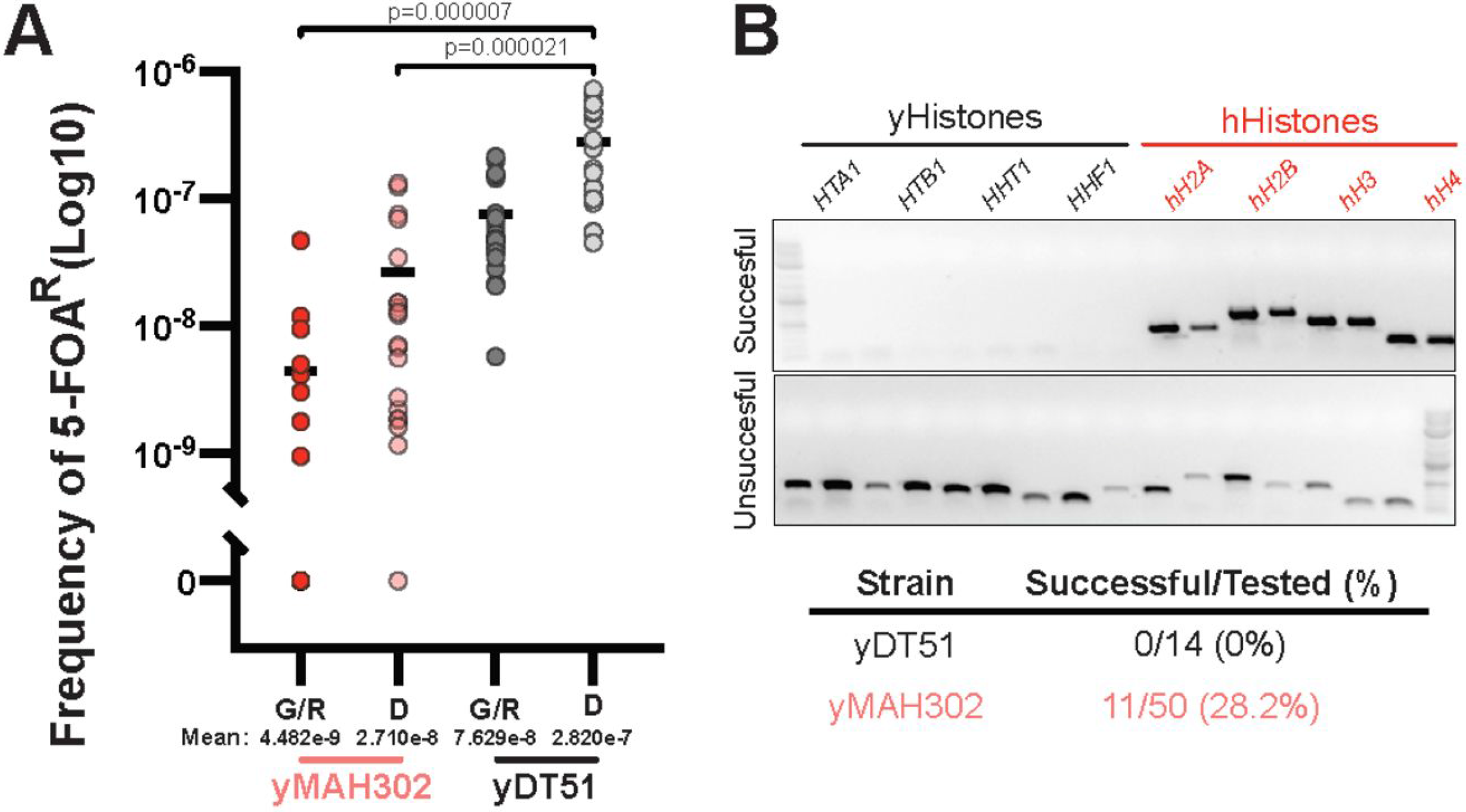
Superloser lowers 5-FOA^R^ mutants and increases isolation of humanized yeast. (**a**) Log10 frequency of 5-FOA^R^ colonies calculated as the number of colonies per plate divided by the total number of cells plated (~1×10^7^ cells per OD600). G/R (Galactose/Raffinose); D (Dextrose). Two-tailed unpaired t-test. (**b**) PCRtag analysis of colonies emerging after 3 weeks of growth. Top, example PCRs from either a successful humanization event and an unsuccessful event, as indicated by the retention of the yeast histone genes. Bottom, summary results from the PCRtag analysis.

Unexpectedly, we also observed a decrease in the frequency of 5-FOA colonies in the normal strain on galactose compared to dextrose (Figure 4C, Figure 5A). We are not aware of any mechanisms to explain this decrease in background. However, it is not due to growth rate differences in galactose vs. dextrose media or fewer replication events under galactose conditions because the OD600 of the initial overnight cultures was not significantly different between dextrose and galactose (Figure 6A). Further, no significant correlation between the initial OD600 and the frequency of 5-FOA resistant colonies was found for either yDT51 (R-squared 0.01743; p=0.4167) or yMAH302 (R-squared 2.571e-6; p=0.9934) (Figure 5B) - further supporting that the observed difference in 5-FOA^R^ colonies is not due to growth differences. Lastly, galactose had no effects on plasmid retention in our other experiments. Therefore, we suggest that galactose seems to have a net negative effect on recombination, resulting in fewer overall 5-FOA^R^ colonies, even in the absence of Superloser.

**Figure 6.**
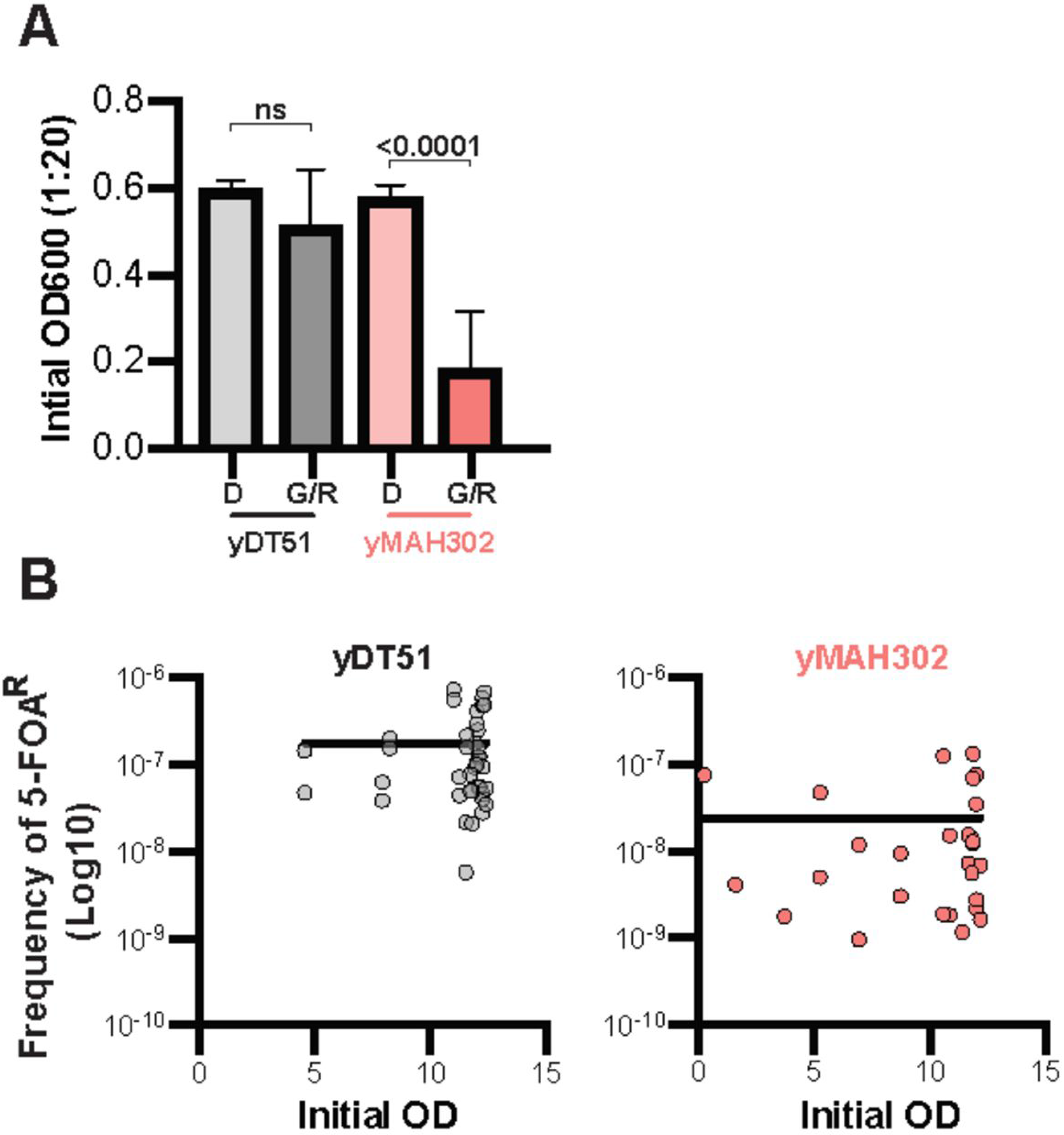
Decrease in 5-FOA^R^ in yDT51 from galactose is unrelated to growth rate differences. (**a**) The initial measurement of OD600 prior to plating the culture onto SC-Trp+5-FOA plates, note that galactose had no effect on the mean initial OD600 of yDT51 (n=20); two-tailed unpaired t-test. (**b**) Plots of initial OD versus calculated frequency of 5-FOA resistance. Semi-log nonlinear regressions are shown.

## Discussion

Humanization of entire pathways or complexes in yeast is a largely unexplored frontier with few exemplars (Wildt and Gerngross 2005; Hanly *et al.* 2014; Agmon *et al.* 2017; Truong and Boeke 2017). Better tools are therefore needed to ensure the ease with which essential yeast genes can be humanized. A major limitation in this is successfully replacing the yeast genes with the human, as the human homolog frequently results in a net decrease in fitness. Because of this, any individual cells that maintain the yeast homolog have a huge fitness advantage. We set out to design a plasmid to minimize this background in plasmid shuffling, especially those with a large fitness cost. We designed in three features to this plasmid that we reasoned would substantially lower the background - strategically reduced sequence identity, two sequence-orthogonal *URA3* markers, and use of an inducible system to remove the plasmid. We demonstrated the utility of the Superloser plasmid by shuffling histones from yeast to human. In the original histone humanization experiments, a total of 7 successfully humanized clones were isolated (Truong and Boeke 2017). Combining the data from the initial study and the work presented here we calculated the frequency of observing a true humanization event as the number of true positives divided by the total number of cells plated across all experiments. Doing so gives a frequency of successful humanization of 1.125e-8 for the conventional shuffle vector and a frequency of 1.47e-7 for the Superloser shuffle vector - a 13.1-fold increase in observing this rare event. Based on the data presented here all three features work together to significantly reduce the background of plasmid shuffling experiments and thereby improve the chances of isolating true positive events.

The Superloser system is likely to be extremely useful for almost any plasmid shuffling application where background is a problem, such as the selection of rare mutants eliciting a highly specific, but rare phenotype, such as e.g. osmotic remediability, in an essential gene. One important and timely potential use of this plasmid lies in the area of complex pathway engineering in yeast, especially considering increasingly large and ongoing genome construction efforts (Annaluru *et al.* 2014; Mitchell *et al.* 2017; Richardson *et al.* 2017; Shen *et al.* 2017; Wu *et al.* 2017; Xie *et al.* 2017; Zhang *et al.* 2017). We imagine that this tool could be used for the humanization (or from other species) of other essential genome architectural proteins such as cohesins, histone modifying complexes or chromatin remodelers, additional histone variants, as well as metabolic pathways. Further, this tool is not limited only to the study of orthologous proteins, but can be readily used for the study of proteins with no identified orthologs or cases in which the orthologs do not complement This tool also offers new opportunities for introducing entirely synthetic pathways in place of endogenous ones to one day confer entirely new functions in budding yeast. In summary, we conclude that Superloser is an extremely useful shuffling vector for use in *S. cerevisiae*.

## Acknowledgments

We thank Leslie Mitchell for the suggesting the name Superloser during lab meeting, Michael J. Shen for help with counting colonies, and members of the Boeke lab for helpful critiques and comments on the data shown here. This work was supported by NIH (NIGMS) postdoctoral fellowship F32GM116411 to D.M.T. and a Genome Integrity Training Program NIH (NIGMS) T32GM115313 to M.A.B.H.

J.D.B. is a founder and Director of the following: Neochromosome, Inc., the Center of Excellence for Engineering Biology, and CDI Labs, Inc. and serves on the Scientific Advisory Board of the following: Modern Meadow, Inc., and Sample6, Inc.

